# Homing and Egg Discrimination in the Western Slimy Salamander, *Plethodon albagula* (Caudata: Plethodontidae)

**DOI:** 10.1101/388249

**Authors:** Robyn R. Jordan, Joseph R. Milanovich, Malcolm L. Mccallum, Stanley E. Trauth

**Affiliations:** Department of Biological Sciences, Arkansas State University, P.O. Box 599, State University, Arkansas 72467; Department of Biology, Loyola University Chicago, Chicago, Illinois 60660.; School of Agriculture and Applied Sciences, Langston University, Langston, Oklahoma 73050

**Keywords:** Egg Discrimination, Homing, Parental Care, *Plethodon albagula*, Western Slimy Salamander

## Abstract

In some species of vertebrates egg brooding is a costly form of parental care. Therefore, misdirection of parental care can significantly lower a female’s fitness. Because of the maternal investment and increased survivorship to offspring from egg guarding, a brooding female should home to her nest site after being displaced a short distance and discriminate between her own eggs and eggs from other females. In this study, we experimentally tested, in the field, alternative hypotheses concerning homing ability and egg discrimination in a population of nesting western slimy salamanders (*Plethodon albagula*). Fourteen brooding females were displaced 1 m to the left or right of their nest sites (determined randomly) for the homing experiment. Furthermore, brooding females (n = 13) were presented with their own clutches, which were displaced 50 cm to the left or right (determined randomly), and unfamiliar egg clutches at their original nest sites. The females were released at an equal distance from both egg clutches. After 24 hours, 12 displaced females (86%) had returned to their own nest sites and were brooding their egg clutches. Also, after 24 hours, nine test females had returned to their own nest sites and were brooding the unfamiliar egg clutches. No control or test females were present at the other new nest site locations. Therefore, we suggest that brooding female *P. albagula* do home to their nest sites and exhibit indirect egg discrimination.

---

Parental investment theory predicts that parental care can be a significant investment for an individual (Trivers 1972). Parental care, which is any form of parental behavior that may increase the fitness of the parent’s offspring, includes nest preparation, egg production, and supervision of eggs or young (Clutton-Brock 1991). Although parental investment improves offspring survivorship, it can also impact the parent’s ability to produce future offspring (Trivers 1972). In some species of vertebrates, egg brooding is a costly form of parental care (Ng and Wilbur 1995).

Female terrestrial plethodontid salamanders oviposit clutches in moist, protected terrestrial sites where they brood the eggs for several months (Nussbaum 1985). Egg brooding increases egg survival by reducing predation, yolk layering, fungal infection, and desiccation (Forester 1984, Highton and Savage 1961, Synder 1971). Egg guarding, however, can be a costly investment for females. Females may suffer metabolic costs, including injury or death, when defending egg clutches from predators (Jaeger and Forester 1993). Female *Desmognathus ochrophaeus* apparently fast during the brooding period (Forester 1981); whereas, *Plethodon cinereus* feeds opportunistically throughout incubation (Ng and Wilbur 1995). Energy consumption by brooding female while defending egg clutches may reduce her ability to reproduce the following year (Jaeger and Forester 1993). Therefore, misdirection of parental care can significantly lower a female’s fitness (Waldman 1988). This costly misdirection of parental care is avoidable if a female can home to her nest site after being displaced a short distance (Trivers 1972).

Homing behavior includes any movement to a spatially restricted area that is known to the animal (Papi 1992). This includes incidences of homing from short distances after displacement from a nest site and long-term nest site fidelity year after year. Homing to specific sites for reproduction, nutrition, shelter, and hibernation occur in adults of ~50 species of urodeles and anurans (Sinsch 1992). Homing in plethodontids is known from direct observation (Madison 1969) and experimental displacement (Barthalamus and Bellis 1972). The non-migratory *Desmognathus fuscus* returned to its original capture sector after translocations to upstream, downstream, non-stream locations (Barthalmus and Bellis 1972). Many species of *Plethontidae* are known to return to capture sites (Kleeberger and Werner 1982; Jaeger et al. 1993; Madison 1969) including nest sites (Forester 1979; Snyder 1971; Peterson 2000) upon removal.

Different cues or sensory mechanisms are used for homing (Sinsch 1992). Olfactory/chemosensory responses appear to be the major mechanism for *D. fuscus, P. jordani*, (Barthalmus and Bellis 1972; Madison 1969; 1972), but not in *P. cinereus* (Jaeger et al. 1993). The sensory mechanisms required for clutch recognition is an important aspect of homing in plethodontids allowing females to distinguish their eggs from unattended clutches (Forester 1979). However, plethodontid females may brood more developmentally-advanced clutches of conspecifics instead of their own. Olfaction was essential for clutch recognition by *D. ochrophaeus*, but is supported by visual and tactile reinforcement (Forester 1979). Displaced brooding female *P. cinereus* also recognized their nest site location by its structure and habitat features (landmarks) (Peterson 2000). Species whose egg clutches are highly susceptible to displacement from the nesting site, such as *D. ochrophaeus*, are under higher selective pressure to evolve clutch and egg discrimination and nest site recognition (Forester 1977; 1979). Terrestrial plethodontids such as *Plethodon albagula* undergo direct development in a terrestrial setting guarded by brooding females (Pough et al. 1998; Trauth et al. 2006; Trauth et al. 2004) and thus have a lower chance for clutch displacement.

Behavioral studies involving homing and egg determination in plethodontids have typically focused on members of *Desmognathus* and smaller bodied *Plethodon*, chiefly *P. cinereus*. Despite their abundance and diversity within woodlands (Burton and Likens 1975a), Hocking and Babbitt 2014; Milanovich and Peterman 2016; Semlitsch et al. 2014), importance to forest ecosystems (Davic and Welsh 2004), and differences in life history traits from smaller bodied *Plethodon*, little information exists about homing and egg discrimination in larger bodied *Plethodon* (see Mathis 1995 for review), especially those of the widespread *Plethodon glutinosus* (slimy salamander) species complex. The purpose of this study was to test alternative hypotheses concerning homing ability and egg discrimination in an aggregation of nesting females of *P. albagula*, to fill in missing gaps in our understanding of parental investment in this species. We hypothesized that female *P. albagula* can recognize their eggs and home to their nests when removed. We predicted that control females would remain at their own nest sites, and displaced females would return (or home) to their own nest sites, and that females will be able to discern between their eggs and those of other individuals.

## Materials and Methods

*Plethodon albagula* was the subject of this study. It is a large woodland salamander found in the Interior Highlands of Arkansas, Missouri and Oklahoma, and parts of Texas (Conant and Collins 1998; Trauth et al. 2004; Highton et al. 1989). Female *P. albagula* brood their eggs cryptically under rocks, logs, or in underground nesting chambers where brooding behavior cannot normally be directly observed. However, a unique opportunity to observe brooding females exists at an abandoned mine shaft in Garland County, Arkansas (Trauth et al. 2004; Trauth et al. 2006; Ford 2008). The shaft is a straight tunnel, approximately 2 m high and 1.5 m wide, extending approximately 149 m horizontally into a hillside, and was excavated in 1880 and abandoned by 1890. Female *P. albagula* migrate into the mine during late summer (July-August), oviposit in August–October, and brood their eggs until hatching in December. Females and young often overwinter in the mine gradually dispersing into the environment December – January. Females lay their clutches in depressions and cavities in the shaft wall formed when the mine was dug. Most of the nest sites are readily visible and accessible to humans. Because females can be observed without disturbance before, after and throughout the brooding period, this site provides a unique opportunity for studying the reproductive ecology of *P. albagula*.

**Figure 1.**
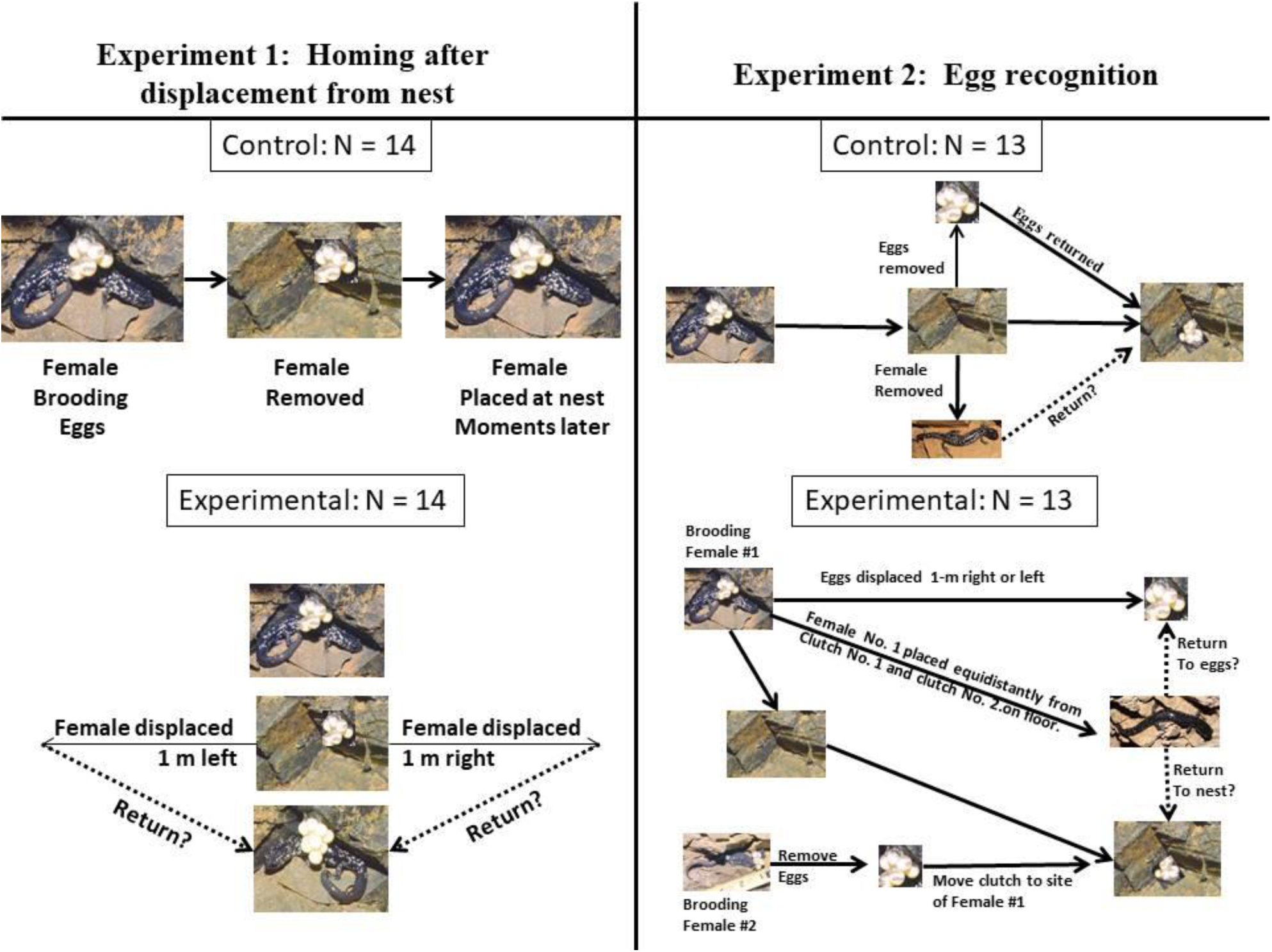
Experimental design for testing homing (left) and egg recognition (right) by female Western Slimy Salamanders (*Plethodon albagula*) in a field setting.

During the first year of the field study (16–17 November 2001), we tested the homing ability of females with a manipulative experiment following methods of Peterson (2000). We randomly selected 28 females and randomly assigned them to a control (*n* = 14) and experimental group. Control animals were removed from, and then returned to, their egg clutch to control for clutch abandonment resulting from handling and or nest disturbance (Fig. 1). Experimental females were randomly displaced 1 m to the left or right of the nest site. We returned to each nest site 24 hours later to determine if females returned or abandoned their nests. Logistic regression was used to test significance (ά = 0.05) between the experimental and control responses.

In year two, (2–3 November 2002) we tested for brooding females’ ability to recognize their own eggs following Peterson (2000). We randomly selected 39 females, and assigned them to a control (*n* = 13) and two experimental groups. The control animals were removed from their nests, and released on the floor directly below their nests (Fig. 1). Their clutches were removed from their egg stalks and immediately returned in the original nest site. Thirteen pairs of experimental females with similar sized egg clutches (± 2 eggs), egg diameter (± 0.64 mm), and developmental stage were selected at random.

We removed an experimental pair of females from their nest sites and placed them in Gladware housing chambers lined with damp filter paper. Then we removed the females’ clutches from their egg stalks. Egg clutches from the first experimental group were displaced randomly 1 m to the left or right of the original nest site location. Egg clutches in the second experimental group were placed at the nest site of the paired member of the first experimental group. The test female was released from the shaft floor at an equal distance from both egg clutches. After 24 hours, we examined each nest site and recorded whether females were present or absent, and whether nest predation had occurred. After the experiment, we returned the clutches to their original positions and released the non-experimental females back at their original nest sites. Logistic regression was used to test significance (ά = 0.05) between treatments.

## Results

In 2001, most displaced females (11/13 [85.7%]) returned to, and control females (92.9% [12/13]) remained with, their own nest sites 24 hours post-handling. There was no significant difference between the two groups (G = 0.383, df = 1, P = 0.536).

In 2002, after 24 hr, most experimental (9/13) and control (12/13) females returned to their own nest sites. There was no significant difference between the groups (G = 2.357, df = 1, P = 0.125). All experimental returnees were observed brooding the opposing salamander’s egg clutches. Neither control nor experimental females moved to egg clutches that were displaced from the nest site There was no significant difference between the number of control and experimental females present at their own nest sites versus present at other nest sites (G= 2.357, df = 1, P = 0.125).

## Discussion

Parental investment theory predicts a significant investment for females that care for their young (Trivers 1972). Fitness can be significantly lowered due to misdirected parental care. Therefore, evolution of a mechanism for identifying one’s own egg clutch is assumed to avoid these fitness costs (Trivers 1972).

This species appears to use the nest site as a proxy for egg clutch recognition. This mechanism for egg recognition likely evolved because in most settings, females are isolated in underground abodes where egg displacement is much less likely due to the nest site’s containment. The environment and number of nest sites in the immediate area can influence the intensity of natural selection on egg discrimination (Peterson 2000). However, the salamanders in our study nested on rock faces, which are more open areas. In this setting, egg clutches are probably dislodged occasionally while fending off nest predation by neighboring females (Ford 2008; Milanovich et al. 2007) and other organisms (Milanovich et al. 2005). On our first visit to the mine, we found an egg clutch at the base of the wall with no female present. Further, members of the *P. glutinosis* complex are known to inhabit (Camp and Jensen 2007) and nest (Gunter 1958; Hines et al. 2004) on rock faces in naturally occurring caves. Observations of *P. albagula* occupation (Briggler and Prather 2006) and nesting in caves also exist, although occupancy appears rare during the winter brooding season (Briggler and Prather 2006). So, there should be selective pressure to recognize displaced egg clutches.

Unfortunately, we did not test the influence of directional displacement, nor did we test if females would return to their empty nest site versus their own displaced clutch. It is possible that the behavioral mechanism for egg recognition requires a behavioral trigger to elicit search behavior for displaced clutches. Further, the most likely place to search for eggs is probably below the nest, rather than to either side. So, even if they had searched, they may have given up and accepted the replacement clutch. In hind sight, it seems probable that return to a vacant nest site may elicit searching as it does in some birds (personal observation). This missing gap in our research would reveal conclusively if egg recognition is restricted to the nest site proxy. However, this site has been closed due to risks of white nose syndrome to resident bats.

Because females were able to relocate their nest sites after displacement, some form of egg discrimination, either direct or indirect, had taking place within this population. Egg discrimination has been documented in *Desmognathus* (Forester 1983) and other plethodontids have homed to specific nest sites [(e.g., *A. aeneus* (Williams and Gordon 1961), *D. auriculatus* (Rose 1966), and *D. ocoee* (Forester 1979)]. Forester (1974; 1979; 1986) and Forester et al. (1983) found female *Desmognathus* were able to locate or home to their own eggs when displaced and when presented with other female’s eggs. However, Peterson (2000) suggested female *P. cinereus* indirectly recognized their own eggs by use of territorial nest sites and suggests the differences between egg recognition abilities may be due to the difference in selective pressures between semi-aquatic and terrestrial plethodontids.

Our data coincide with Peterson (2000) and fail to support the hypothesis that females discriminate their eggs and those of a conspecific. After returning to their original nest sites, test females began to guard the unfamiliar egg clutches. The current study lacks evidence that *P. albagula* can recognize their own clutches. This supports findings with other *Plethodon* sp. where females depend on chemical or environmental cues to locate their nest sites as opposed to their egg clutches after displacement.

## Acknowledgements

We thank D. Saugey for providing an opportunity to examine these mine shaft animals and B. Wheeler, H. Worley for field assistance.

